# Trajectory Recording and Analysis System for Cockroach Robot

**DOI:** 10.1101/2021.11.16.468890

**Authors:** Ruituo Huai, Haoran Zhu, Shuo Yang, Zhihao Yu, Hui Wang, Junqing Yang, Pingqiu Zhang, Yong Shi, Rui Yan

## Abstract

In this study, We design a trajectory recording and analysis system to record and analysis the changes in the movement behavior of the cockroach robot after stimulation. The external hardware of this system is an infrared touchpad as the experimental platform for the cockroach robot to crawl freely, and the infrared matrices densely distributed in the X and Y directions of the infrared touchpad are used to detect and locate the position of the cockroach robot. The cockroach robot’s movement trajectory is displayed visually through the projector’s interface projection on the infrared touchpad. The system software consists of three main parts: the electrical signal parameter setting module, the movement trajectory recording module, and the data analysis module. The electrical signal parameter setting module sets the stimulation parameters and configures the corresponding serial port to independently stimulate the left and right antenna and cercus of the cockroach; the trajectory recording module is used to record the trajectory of the cockroach robot through the coordinate positioning method. The data analysis module explores the change of motion behavior of the cockroach robot with time after receiving the stimulus by using the stage analysis method, and explores the change of motion of the cockroach robot with different voltage stimulus by using the module analysis method. The system is tested in experiments and the results demonstrated its applicability to the recording and analysis of the cockroach robot’s trajectories.

## INTRODUCTION

Since the 1990s, research on animal robots has flourished ^[1–3]^. With their flexibility and stealth ^[4,5]^, cockroach robots are more ideal for research robots because of their potential applications in complex search and rescue and covert detection ^[6–8]^.

Early information about the animal is obtained from the calculations required to control the trackball. The animal was tethered to a freely rotating ball and the ball was brought into direct mechanical contact with a pair of orthogonally positioned rotary encoders, a design that employed an older computer mice or optical sensor to record the trajectory of the insect ^[9–11]^. However, since the trackball is in contact with the encoder, it is not guaranteed to be free of mechanical resistance in all directions of rotation. Immediately, the Kramer-Kugel system eliminates this mechanical resistance and provides detailed information about the paths and speed of the animal. This device is based on a polystyrene ball floating on an air cushion ^[12–15]^. The cricket is loosely tethered to the ball, which rotates under the action of the insect’s motion. The rotation of the ball is detected by a pair of optical encoders that produce variations in the x, y coordinates, which map the path of the ball. The system is designed to measure and record movements over distances of many meters and to generate detailed records of the animal’s behavior in a compact format ^[16]^. Recording insect motion trajectories similarly to hovering balls has also been experimented ^[17–22]^.

Optical patterns reflected from the surface of the trackball are detected simultaneously with sensors for any trackball moving in the forward-backward and left-right directions, and the trackball is designed with a two-dimensional sensor chip to monitor the walking trajectory of crickets ^[23]^. A virtual reality system was proposed based on this research. The system was used to simulate a natural environment in a virtual reality (VR) environmental laboratory to study the movement trajectory of cockroaches exploring behavior in a virtual forest ^[24–28]^. The same approach of hovering balls was used in recording trajectories, which requires tethering the insects to the ball, thus inevitably inducing physical differences in the sensorimotor control of the behavior. On account of the tethering problem, a tetherless trackball recording method has recently been proposed to record the trajectories of ants. This system uses a motion-compensated treadmill combined with a high-speed camera to track ants ^[29]^. However, if the mass of the trackball tied to the animal is more than 1.5 times that of the animal in the process of rotating motion, there will be problems in controlling the inertia of the trackball ^[30,31]^.

The application of sensors has been explored in the study of trajectory recording of insects. Researchers have measured the gait and movement velocity trends of insects by mounting inertial sensors with 6 or 9 degrees of freedom on the animal ^[32,33]^, which were then combined with a camera to record the animal’s trajectory ^[34,35]^. Similar to this method is the recording of insect movement by a 3D motion capture system ^[36–40]^.

In recent years, the study of insect locomotor behavior has been extended to vision. The combination of advanced techniques in neural engineering with a miniature system-on-a-chip containing wireless communication and control circuits has the potential to enable remote and direct control of insect movements. A vision-based automated system has been proposed for the objective assessment of bio-robotic control capabilities in flat arenas and mazes with upright walls ^[41,42]^. This system was designed to identify and detect the position of insects relative to a target line by using Kinect’s infrared depth camera ^[43]^, an approach that requires processing of the visual aspects of the video and may not be as accurate as sensor measurements in recording insect velocity and steering angle. Finally, based on the above discussion, we hope to build a system that not only allows cockroaches to crawl freely but also records the trajectory, speed, turning angle and other information of cockroaches to facilitate us to explore the movement pattern of cockroach robot.

In this paper, the system designed in this paper extends the study to the cockroach robot and maximizes its normal biological instincts. In order to more intuitively observe the real response of cockroach robot under different stimulus parameters. We changed the way the roaches were tethered to allow them to crawl normally. Instead of moving on a trackball, the cockroach moves on an infrared frame. We recorded the real movements of the cockroach robot by collecting the crawling trajectory of the cockroach under different stimulus parameters. As the cockroach robot moves on the infrared frame, different stimuli are applied to it with different parameters and the cockroach robot responds differently. The trajectory of the cockroach robot is recorded by obtaining the X, Y coordinates of the current position of the cockroach robot. This allows obvious observation of the real response of the cockroach robot and a recording of its response to different stimulation parameters.

## MATERIALS AND METHODS

### Animals

These experiments were conducted using colonies of American cockroaches (Periplaneta Americana, length:40-50 mm, width:7-8 mm, weight:5-8 g) raised in our laboratory. Infrared heating lamps and daily spraying of tap water helped maintain temperature and humidity to more closely mimic natural habitat conditions. The colony was fed twice a week, mainly fresh vegetables. Test subjects were selected from the colony, paying attention to their size and activity levels.

### Electrode implantation

Before starting the experiment, the cockroach needs to be operated on and the electrodes are implanted into the antenna and cercus of the cockroach through routine surgery, and the stimulation receiving device connected to it is placed on the back of the animal.

The electrodes are made of stainless steel needles with a diameter of 0.16 mm and a length of 1-5 mm; the connecting wire between the electrodes and the stimulator receiving device is 0.1 mm copper wire; The electrodes were divided into three types according to the implantation site: the electrodes implanted in the antenna of the cockroach were 3-5 mm in length; the electrodes implanted in the cercus of the cockroach were 2-4 mm in length, and the electrodes implanted in the back of the cockroach were 1-3 mm in length.

Before surgery, the cockroaches were placed in an ice-water mixture for 3-5 minutes to anesthesia. The anesthetized cockroach is placed on the cryogenic operating table (see Fig. 1A) for the electrode implantation procedure, and the distal ganglia of the cockroach’s antennae and cercus were trimmed, and two antennal electrodes were implanted to 1 mm from the antennal root, two cercus electrodes were implanted to 0.5 mm from the cercus root, and grounded electrodes were implanted to the back of the cockroach (see Fig. 1B). For each electrode implanted, it was fixed to the corresponding position with 3M glue until all electrodes were implanted. Finally, the postoperative cockroaches were placed under a warm lamp to recover for about 30-60 minutes, and the recovered cockroaches were fed individually as the subjects.

**Fig. 1.**
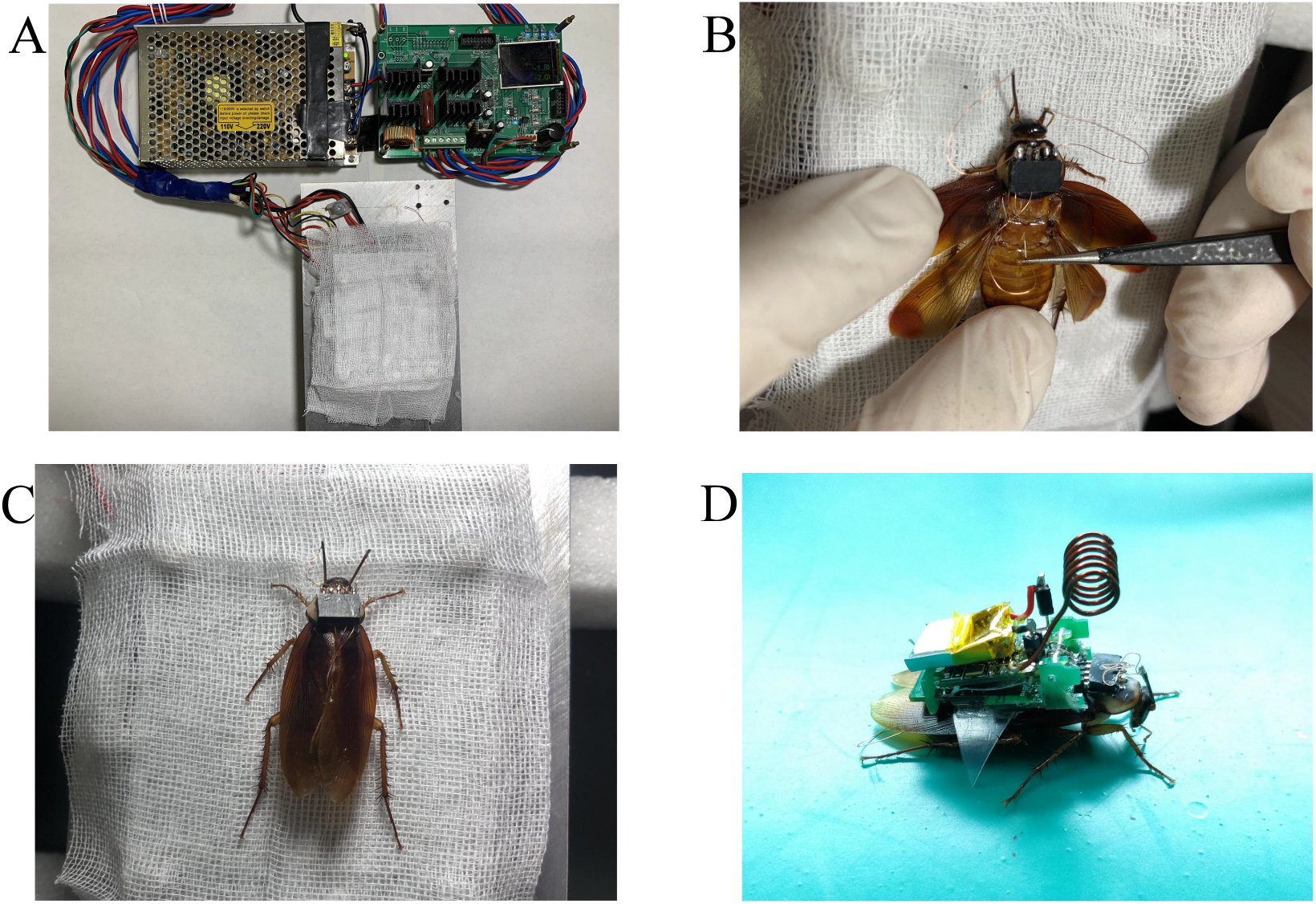
Cockroach surgical equipment and experimental process. (A) The cryogenic operating table is used to keep the cockroach under anesthesia during the cockroach electrode implantation procedure. (B) Cockroach electrode implantation process. (C) Electrode implantation procedure completed. (D) Cockroach with attached electronic backpack (battery on top).

### System external hardware device

The infrared touch panel and projector were selected as external hardware devices for this system. Instead of fixing the cockroach with a tether, the system allows the cockroach to crawl freely on an infrared touchpad with a wireless electronic backpack. The infrared touch panel is not only the experimental platform for the cockroach robot to crawl, but also a carrier for the real-time synchronization of the system software form interface.

The infrared touch panel used in this system is an interactive whiteboard (see Fig. 2A). With the effective reading size of interaction whiteboard is 1811 mm*1296 mm. It Connect to computer through USB interface.

**Fig. 2.**
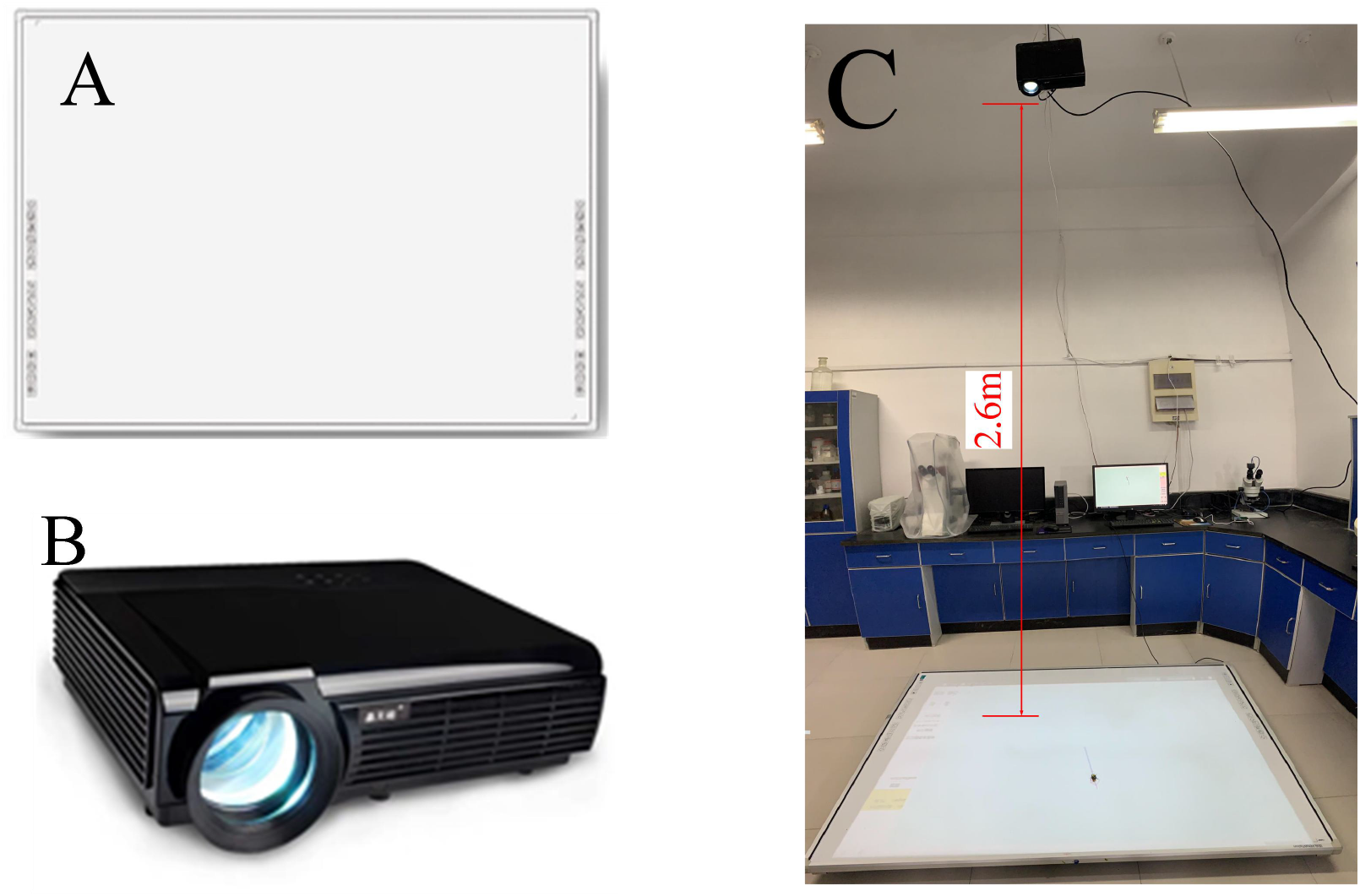
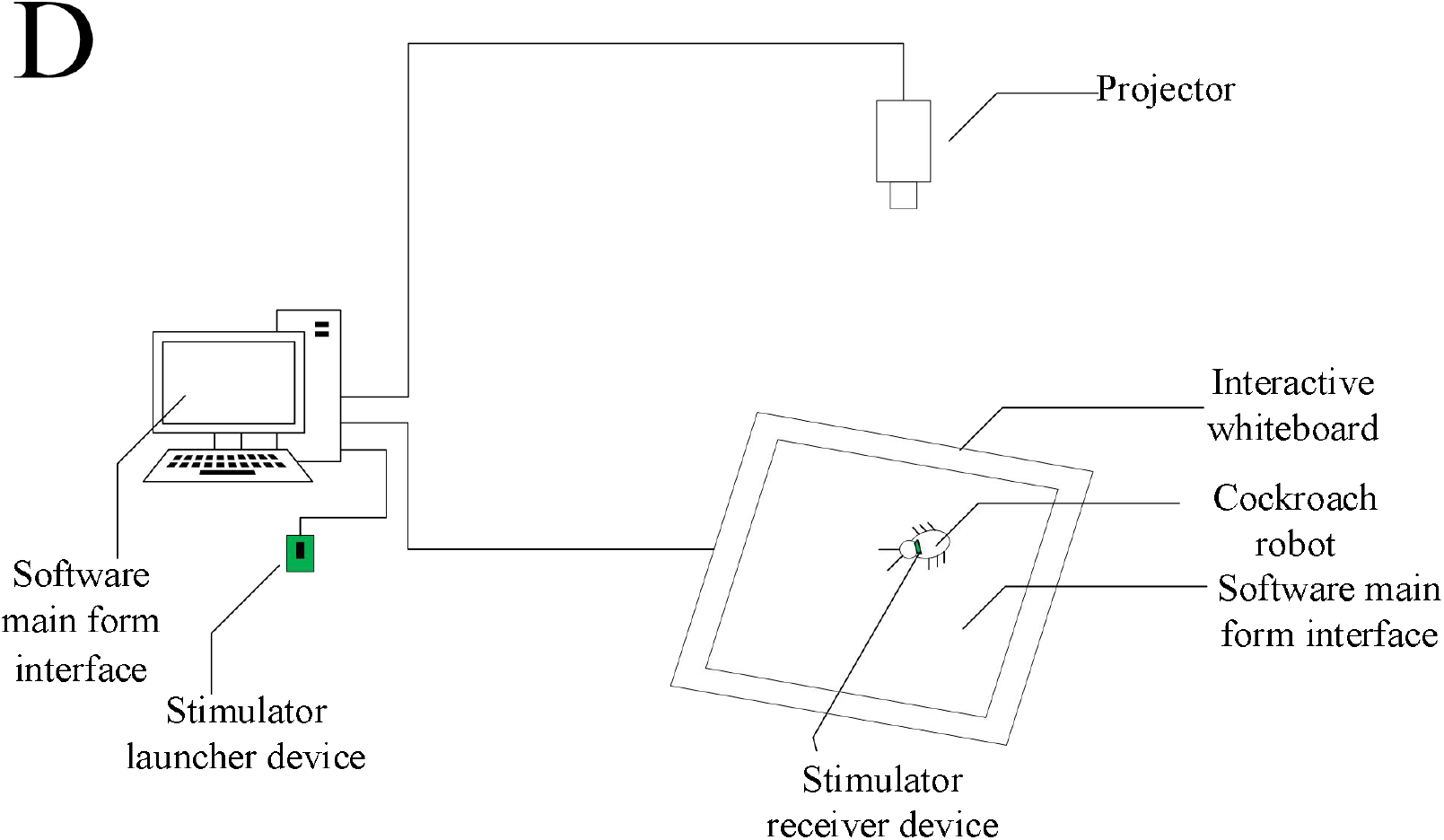
(A) The interactive whiteboard is used to identify the cockroach robot while providing a platform for the cockroach robot to crawl freely. (B) The projector projects the system interface onto the interactive whiteboard, enabling visual observation and display of the cockroach’s movement trajectory. (C) Experimental environment for cockroach robot trajectory recording. (D) Schematic diagram of the overall structure of the system.

The infrared touch panel is the use of the X, Y direction of the dense infrared matrix to detect and locate the location of the touch. It is installed around the outer frame of the circuit board, the circuit board is lined with infrared transmitter tubes and infrared receiver tubes, transmitter tubes, and receiver tubes one by one to form a horizontal and vertical cross infrared array. According to the touch positioning principle of the infrared trackpad, when the cockroach robot crawls on the infrared trackpad, its body will block the infrared array in the X and Y directions of the infrared trackpad, and the position of the cockroach robot can be accurately located by the infrared trackpad. The cockroach robot keeps blocking the infrared array during the crawling process, and the blocked position points together are the movement trajectory of the cockroach robot on the infrared touch panel.

The role of the projector is to project the system software form interface proportionally onto the infrared touch panel. According to the projected image of the projector, the experimenter can observe the location and area of each function module of the main software form located on the infrared touch panel, and can directly conduct the cockroach robot stimulation operation experiment. The projector chosen for the system is a multimedia LCD projector (see Fig. 2B). The projector projection screen size ranges from 60 to 120 inches, and the projection distance is between 1.7 meters and 4.5 meters, which can meet the experimental needs.

### The working process of the system

The overall structure of the cockroach robot motion trajectory recording and analysis system is shown in Fig. 2C, and the principle structure is shown in Fig. 2D.

System experimental process is as follows:

1. Firstly, open the software form interface of the motion track recording and analysis system on the PC. Then synchronize this window interface to the infrared touchpad. Finally, debug and control the experimental system equipment until the entire window interface of the Motion Track Recording and Analysis System software is displayed in real-time on the infrared touchpad.
2. Set the stimulation parameters for each stimulation site on the PC, attach the stimulator receiver device to the back of the cockroach robot, and place the cockroach robot in the track recording module window area on the infrared touch panel where the system software window interface has been projected.
3. Pressing the stimulation button on the infrared touch panel, the stimulator transmitter sends the stimulation signal to the stimulator receiver through the wireless communication module; the stimulator receiver converts the stimulation signal into the corresponding micro-current applied to the tentacles and tail whiskers of the cockroach robot; during the movement of the cockroach robot, the motion track recording module records the entire motion track and data information of the cockroach robot in real-time.
4. Finally, the recorded and saved motion data is statistically analyzed in the data analysis module.

### System software structure and running results

The system software is mainly composed of three functional modules: the electrical signal parameter setting module, the motion trajectory recording module, and the data analysis module. The main window interface of the software (see Fig. 3A) contains three main modules and several small functional modules, such as the playback module, the stimulus data display module, and the selected start recording module. The software main form interface is to integrate the functions of each module, divide the position interval of each functional module, and become the main body of the interface synchronization of the infrared touchpad, as the main experimental area of the cockroach robot.

**Fig. 3.**
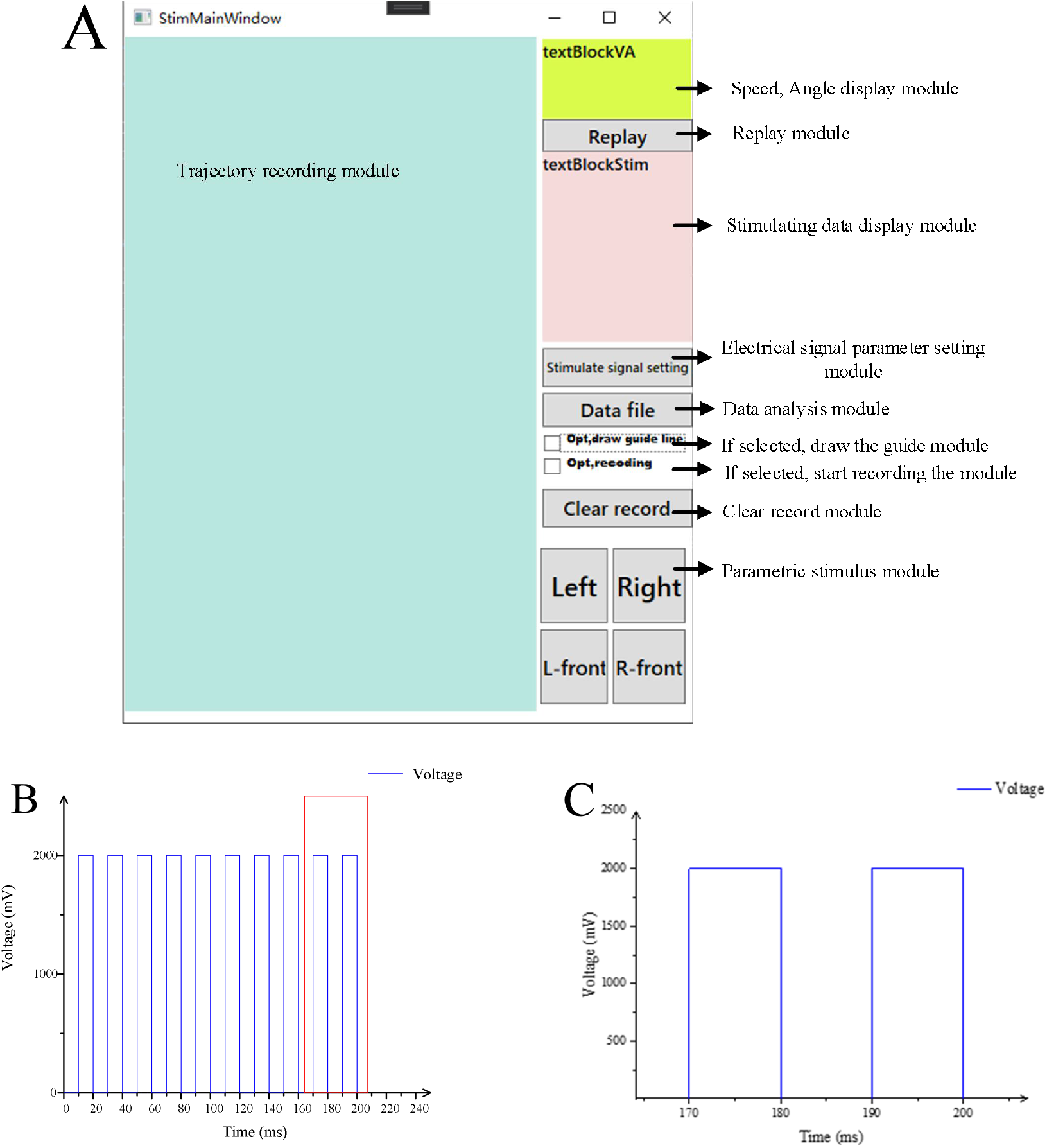
Software main form interface and parameter waveform display. (A) Software main form interface. (B)The stimulation waveform is generated by the electrical signal parameter setting module. (C) Local magnification of the stimulation waveform.

The parameter setting module of the electrical signal is the basis for the system to stimulate the cockroach robot. The function of this module is to set various experimental parameters required to stimulate cockroaches and then send the set parameters to the stimulator launcher through the serial port. It then communicates with the stimulator receiver to generate a corresponding micro-current to stimulate the robot cockroach. The four buttons of parameter stimulation left, right, left front and right front correspond to the four stimulation positions of antenna and cercus on the left and right sides of the cockroach.

The motion trajectory recording module can record the real motion path and trajectory data of the cockroach robot in real-time, with different markers to distinguish the motion direction and stimulation position. The cockroach robot’s current position on the infrared touchpad is marked by a red diamond-shaped arrow and a blue straight line. The diamond indicates the current position of the cockroach robot on the infrared touchpad and the direction of the arrow indicates the positive direction of the robot’s current movement; the blue line is located at the bottom of the red arrow and is twice as long as the red arrow, to clearly show the positive direction of the cockroach robot’s current movement. The stimulus point marker consists of a white circle, a green arrow, and a blue arrow. The position of the white circle indicates the stimulus position of the cockroach robot when it moves to that point; the direction of the green arrow at the bottom of the circle indicates the positive direction of the cockroach robot’s movement at the moment the stimulus occurs at that position point. The direction of the blue arrow on the outside of the circle indicates the stimulus applied to the cockroach robot at that location point (see Fig. 4.).

**Fig. 4.**
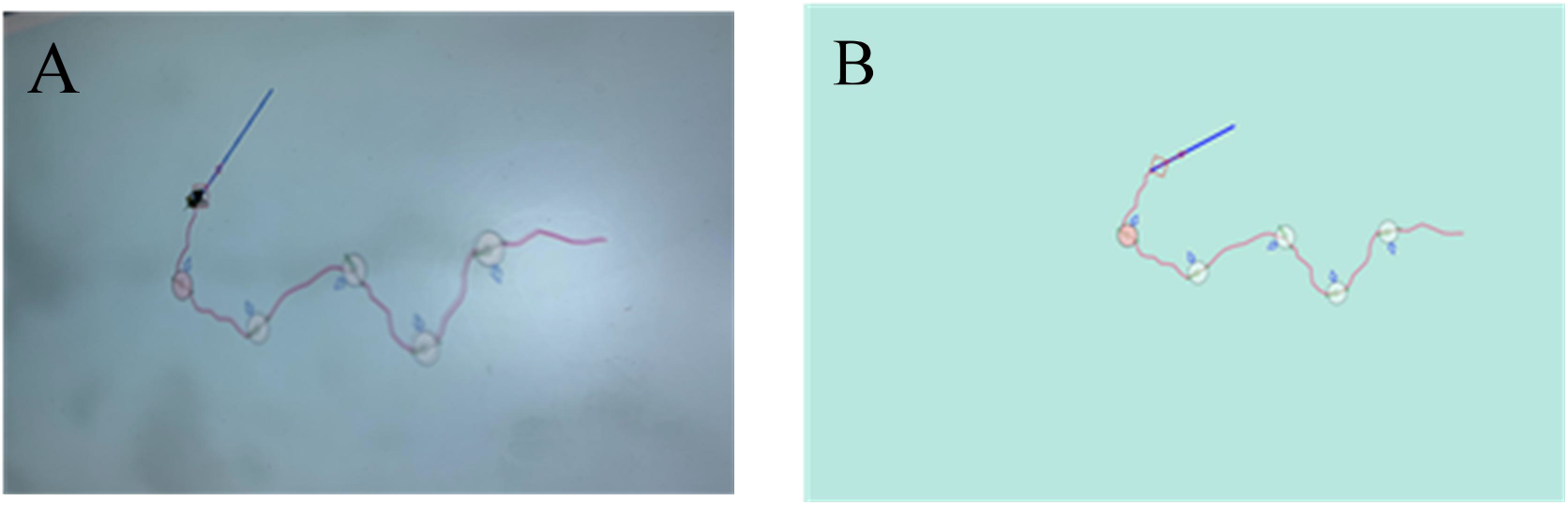
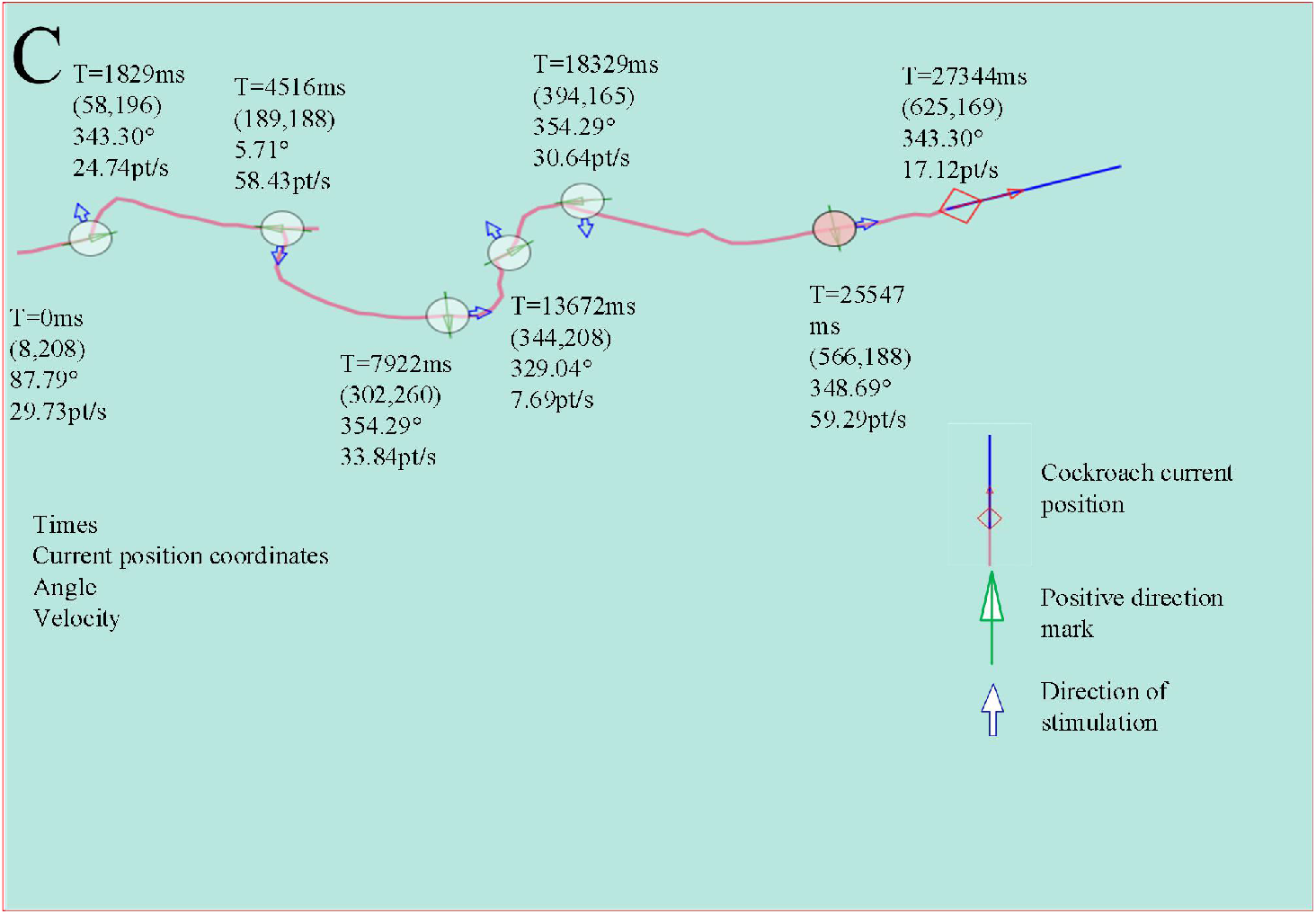
Actual crawling trajectory of cockroach robot. (A) Infrared touchpad to capture the movement trajectory of the cockroach robot. (B) The system saves the movement trajectory of the cockroach robot. (C) The system records the complete cockroach movement trajectory diagram, which response to the location and time of stimulation of the cockroach, and records the movement trajectory, turning angle and movement speed of the cockroach.

The motion trajectory recording module is used to record the motion of the cockroach robot by obtaining the coordinates of the position of the cockroach robot on the main window of the infrared touchpad. Real-time analysis of each new motion position of the cockroach robot is the key to motion track recording. The system is designed to resolve new positions in four steps.

#### (1) Data initialization

Before the experiment is conducted, the data information such as position coordinates, time, angle and speed are initialized to zero.

#### (2) The determination method of new position appearance

Every time a new position is moved, the size of the distance between the new position point and the old position is to be judged, and moving too small is not analyzed. The system sets the minimum distance of the moving position as 10 pt. When the change of X and Y coordinates of the new and old position points is less than 10 pt at the same time, the movement is considered too small and the new position point is not judged.

#### (3) New position speed and angle calculation

The velocity of the new position is the average velocity calculated from the ratio of the distance between the old and new position points to the time. The velocity of the cockroach robot’s new position motion is calculated as:

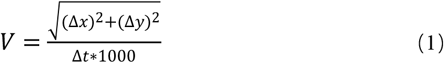

represents the speed at which the cockroach robot moves to the new position, the change in X coordinate between the old and new points, the change in Y coordinate between the old and new points and the time interval between the old and new points in ms.

The angle of the new position is derived from the change in coordinates of the old and new positions in radians, and then the radian value is converted into an angle. The cockroach robot rotation angle is calculated as:

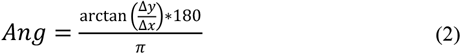

The angle of rotation between the old and new positions of the cockroach robot is represented as an inverse tangent function. If the calculated angle result is less than 0, 360° will be added to the calculated result as the movement angle of the current position; if it is less than 0, 180° will be added to the calculated result before the movement angle of the current position.

#### (4) Set the new position point as the old position point

After the new position is resolved, the position data is saved to become the old position point, and then we wait for the new position to appear.

The above cycle is repeated for each new position point, new position points are connected and the track is displayed in real time in the track record sub-module form interface.

The actual crawling trajectory of the cockroach robot under the stimulation effect is shown in Fig. 4.A, taking the stimulation parameter voltage amplitude of 1500mV, signal period of 10ms, pulse repetition period of 20ms and number of pulses of 10 as examples. The path saved on the PC side is shown in Fig. 4B. By comparing the actual path map on the infrared touchpad with the path map saved on the PC side, the paths of the motion trajectories in the two maps are the same.

During the experiment, we stimulated the cockroach robot with different stimulation parameters to change its movement direction according to the actual situation. The motion trajectory recording module enables us to see clearly the trajectory traveled by the cockroach robot, as well as information on the location of stimulation, the direction of stimulation, the positive direction of cockroach movement at the time of stimulation, and the current position. After the experiment, the data recorded by the data analysis module was combined to analyze the movement of the cockroach robot (see Fig. 4.C). The data analysis module recorded (see Table 1.) eleven types of data information about the motion of the cockroach robot on the infrared touch panel, namely: data line number (ID), stimulus type (TYPE), time (time, ms), X coordinate (pos X), Y coordinate (pos Y), voltage amplitude (Amp, mV), signal period (T), pulse repetition period (Tw), number of pulses in a group (N), angle change(Angle, °), velocity change(Veloc, pt s^−1^). Where NONE_STIM stands for no stimulus, DERCTION_LEFT stands for left stimulus, DERCTION_LEFT_FORWARD stands for left front stimulus, and DERCTION_RIGHT stands for right stimulus.

**Table 1.** The data recording module records the movement data of cockroaches

| ID | TYPE | Time | Pos X | Pos Y | Amp | T | Tw | N | Angle | Veloc |
| --- | --- | --- | --- | --- | --- | --- | --- | --- | --- | --- |
| 0 | NONE_STIM | 0 | 8 | 208 | 0 | 0 | 0 | 0 | 87.79 | 29.73 |
| 101 | DIRECTIN_LEFT | 1829 | 58 | 196 | 2400 | 10 | 20 | 25 | 343.30 | 24.74 |
| 252 | DIRECTION_RIGHT | 4516 | 189 | 188 | 2400 | 10 | 20 | 25 | 5.71 | 58.43 |
| 448 | DIRECTION_LEFT_FORWARD | 7922 | 302 | 160 | 2800 | 10 | 20 | 25 | 354.29 | 33.84 |
| 770 | DIRECTION_LEFT | 13672 | 344 | 208 | 2400 | 10 | 20 | 25 | 329.04 | 7.69 |
| 1033 | DERCTION_RIGHT | 18329 | 394 | 165 | 2400 | 10 | 20 | 25 | 354.29 | 30.64 |
| 1420 | DIRECTION_LEFT_FORWARD | 25547 | 566 | 188 | 2800 | 10 | 20 | 25 | 348.69 | 59.29 |
| 1522 | NONE_STIM | 27344 | 652 | 169 | 0 | 0 | 0 | 0 | 343.30 | 17.12 |

### Data analysis methods and Result

The current speed and position angle of the cockroach robot are calculated in the motion trajectory recording module. To calculate the change in speed and angle of the cockroach robot’s motion after stimulation, we must first determine the change in the cockroach robot’s motion during the time interval from the occurrence of the stimulus to the end of the response. Cockroach’s antenna and cercus have different reaction times to stimulation, so the system treats the three types of data separately: forward, left turn and right turn. The system divided the response of the cockroach robot’s forward speed and rotation Angle with time from 0 to 1200ms into 12-time intervals every 100ms for statistical analysis.

In the experiment of forward stimulation of the cockroach robot, the robot’s movement speed changes variously under different voltage stimulation of the cockroach. Therefore, the algorithm is designed in two steps: (1) The mean value of the velocity of the cockroach robot in a certain time interval is subtracted from the current instantaneous velocity of the cockroach robot at the time of stimulation. (2) Take the positive part of the difference to be the change in the speed of the cockroach robot. Taking the 0-100ms time interval as an example, the sum of the velocities of all the experimental data points of the change in motion of the cockroach robot after stimulation can be expressed as:

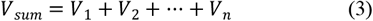

The mean value of the change in velocity of the cockroach robot over the 0-100ms time interval can be expressed as:

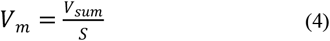

Where S is the number of all experimental data points of the cockroach robot within 0-100ms.

The rotation angle is calculated concerning the position of the cockroach robot at the time the rotation occurs after the cockroach robot has been stimulated to turn left and right. The angle recording period in the motion trajectory recording module is 360° and is divided into four quadrants. When the cockroach robot is moving left, it is rotating counterclockwise around the zero point (see Fig. 5A), while the angle recording in the system’s trajectory recording module is incremental in a clockwise direction (see Fig. 5.B). Therefore, in designing the algorithm for calculating the left rotation angle of the cockroach robot, the system divides the position interval where the cockroach robot’s stimulus occurs into four parts: 0°-90° first quadrant, 90°-180° second quadrant, 180°-270° third quadrant and 270°-360° fourth quadrant.

**Fig. 5.**
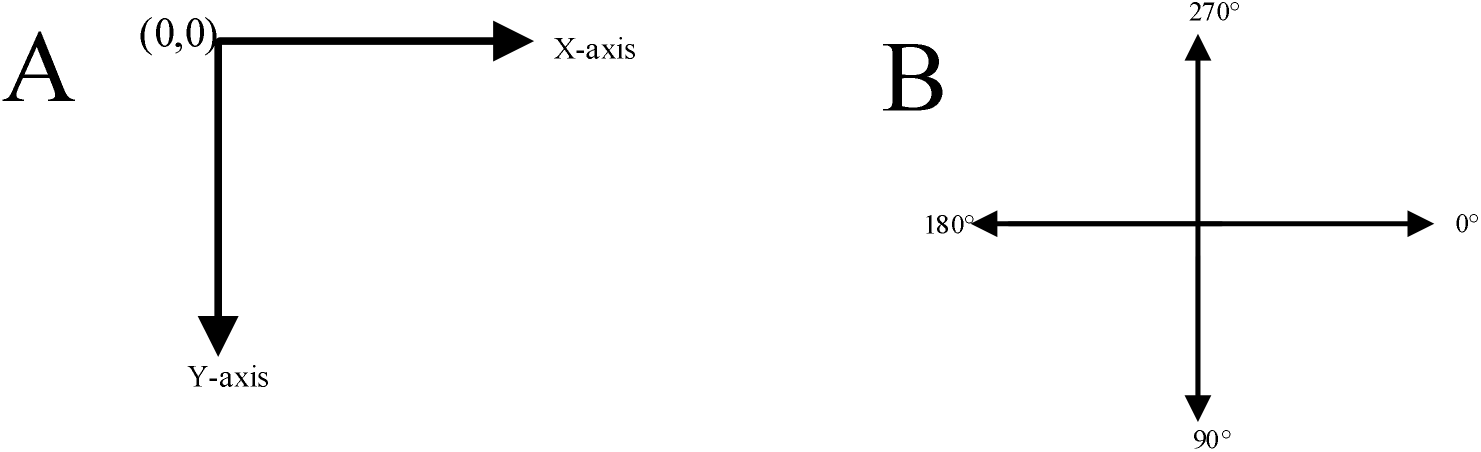
Schematic diagram of coordinate axis and angle direction of track recording module. (A) Main form interface coordinate axis direction. (B) Main form interface angle direction.

In the fourth quadrant, for example, if the difference between the mean value of the cockroach robot’s position angle in the 0-100ms period and the angle of the cockroach robot’s position at the time of stimulation is negative, then the actual rotation angle of the cockroach robot should be the absolute value of this difference. The specific calculation can be expressed as follows:

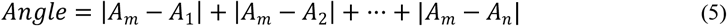

If the difference between the mean value of the cockroach robot position angle in the 0-100ms period and the angle at which the cockroach robot is positioned at the time of stimulation is positive, then the actual rotation angle of the cockroach robot should be 360° minus this difference. The specific calculation can be expressed as follows:

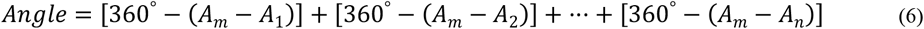

The right turn stimulus algorithm design is similar in principle to the left turn stimulus algorithm design and opposite in outcome, the difference being that the left turn is a counter-clockwise rotation of the quadrant of the track recording module and the right turn is a clockwise rotation. This opposite motion should also be reflected in the algorithm. The actual angle of rotation of the cockroach robot is the difference between the mean value of the cockroach robot’s position angle in a given time interval and the angle at which the cockroach robot was positioned at the time of stimulation is a positive value. The specific calculation can be expressed as follows:

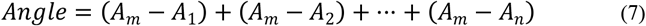

If the difference between the mean value of the cockroach robot position angle and the angle at which the cockroach robot is positioned at the time of stimulation is negative, then the actual rotation angle of the cockroach robot should be 360° plus this difference. The specific calculation can be expressed as follows:

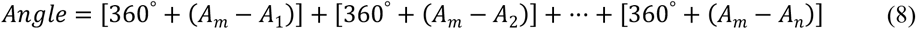

*Angle* is the rotation angle of the cockroach robot during the 0-100 ms time interval, Am is the average value of the angle swept by the cockroach robot during the 0-100 ms time interval, *A*_1_, *A*_2_, ⋯, *A_n_* represents the angle swept by the cockroach robot during the rotation.

In the experimental process, taking 2 V forward stimulation as an example, the system conducted stimulation experiments on multiple cockroaches, and the data points for each 100 ms time interval after the 0-1200 ms time of stimulation were statistically analyzed, and the results are shown in Fig. 6. From the results, it can be seen that the change in speed and rotation angle of the cockroach robot was small during the 0-300 ms time interval, and the movement speed and rotation angle of the cockroach robot changed significantly during the 300-400 ms time interval the movement change of the cockroach robot stabilized after 800-900 ms.

**Fig. 6.**
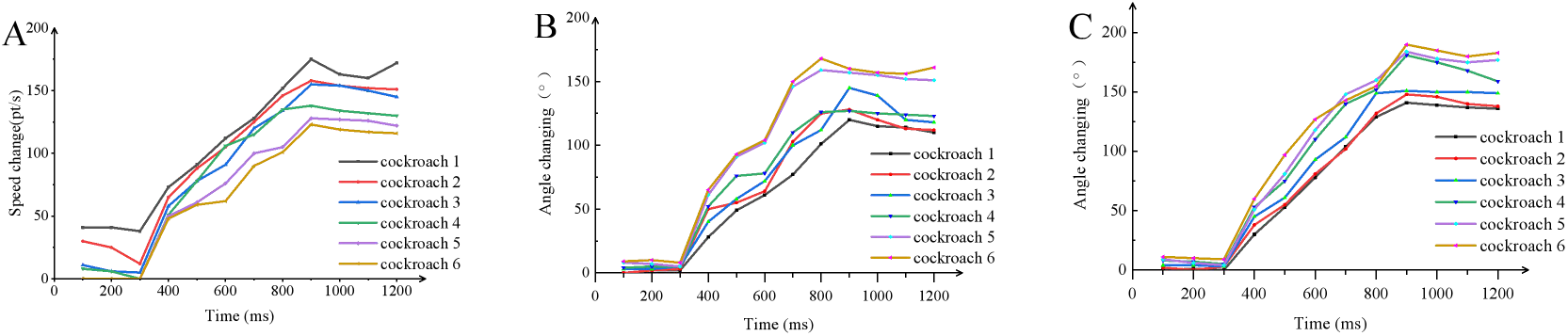
Results of the response over time after stimulation of the cockroach robot (A) Response results of forward speed of six cockroach robots changing with time. (B) Response results of left turning angle of six cockroach robots changing with time. (C) Response results of right turning angle of six cockroach robots changing with time.

The stimulus parameter that has the greatest influence on the cockroach robot during the real experiments is the voltage amplitude. The algorithm design of the response law with stimulation voltage is based on the temporal law algorithm supplemented with the changes in the forward speed and rotation angle of the cockroach robot at voltages of 1000mV, 1500mV, 2000mV, 2500mV, and 3000mV. Based on the results of the cockroach robot’s response to changes with time after stimulation, the algorithm uses a time interval of 800-900ms for the end velocity mean *V_m1_* and the end angle mean *A_m1_*.

In the implementation of the voltage variation response law algorithm. The cockroach robot forward speed with stimulus voltage change response law algorithm is divided into five steps: (1) Deserialize the XML data document. (2) Traversal the set of row data to determine the stimulation method. (3) Calculate the average value of velocity in 800-900ms time interval according to the stimulation method. (4) Determine whether the stimulation voltage is within the five-module interval based on the voltage amplitude in the row data set. (5) Calculate the difference between the mean velocity value and the initial velocity within 800-900ms. The result is the increase in velocity of the cockroach robot at that stimulation voltage (see Fig. **7.**A).

**Fig. 7.**
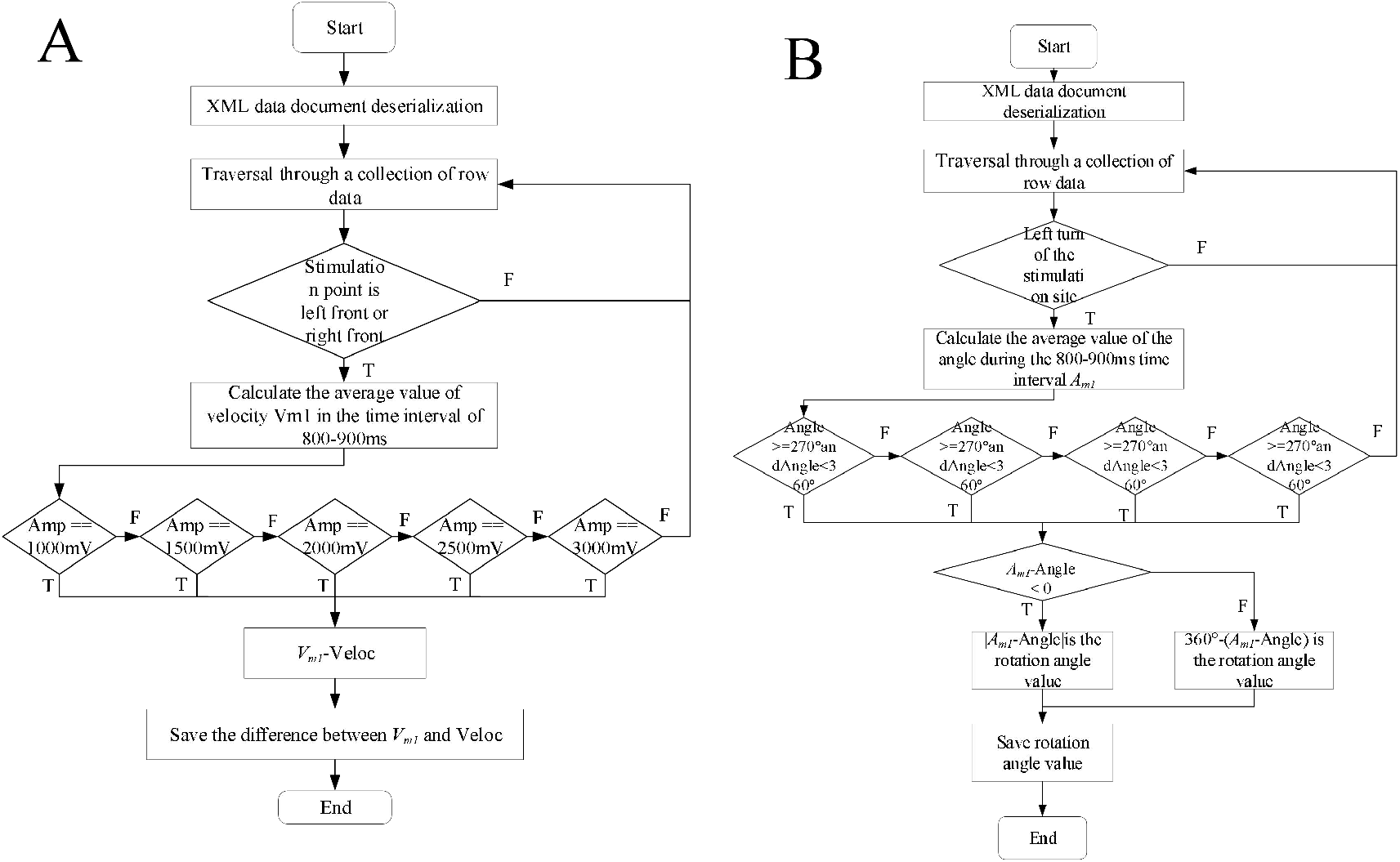
(A) Algorithm design flow chart of cockroach robot’s forward speed changing with stimulation voltage. (B) Algorithm design flow chart of cockroach robot’s left turn angle changing with stimulation voltage.

The left rotation angle calculation algorithm is similar to the forward speed calculation algorithm, with the first two steps being the same and the following steps: (1) Calculate the average value of the angle during the 800-900ms time interval. (2) Determine in which interval the initial angle of the cockroach robot before rotation in the row data set is located. (3) Determine whether the difference between the mean angle value and the initial angle in the 800-900ms time interval is less than zero. When the result is less than zero, the absolute value of the difference between the mean value of the angle and the initial angle in the 800-900ms time interval is the value of the rotation angle of the cockroach robot under this stimulation voltage. When the result is greater than zero, the difference between 360° and this result is the value of the cockroach robot rotation angle under this stimulation voltage. The right-turn stimulation algorithm is designed in the same way as the left-turn stimulation algorithm (see Fig. **7.**B).

The results of the statistical analysis of the response of the cockroach robot movement to voltage changes are shown in Fig. 8. The results of the forward left and right rotation movements of the cockroach robot under different voltage stimulation show that the forward speed and rotation angle of the cockroach robot increase with the increase of voltage in the range of 1-3 V.

**Fig. 8.**
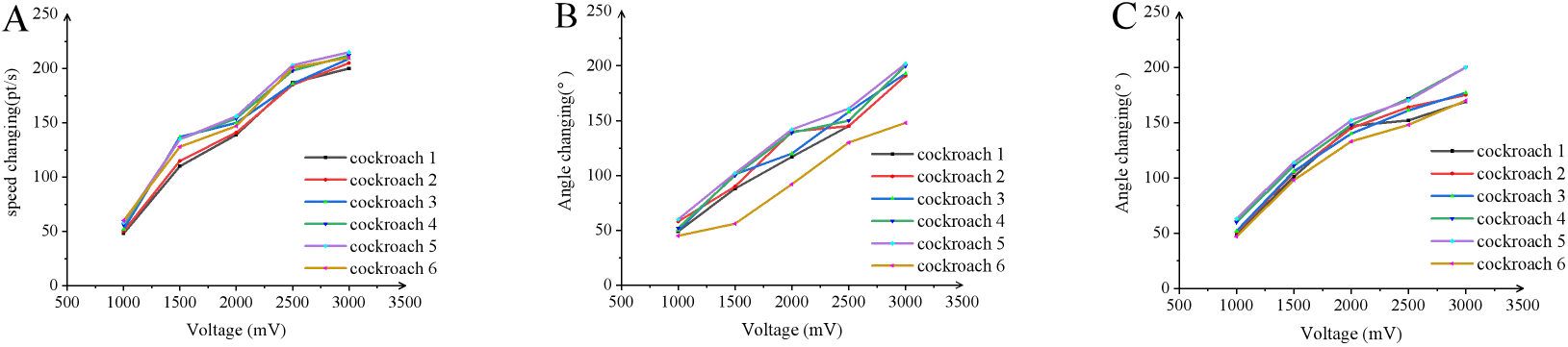
Reaction result of cockroach robot with voltage change. (A) Response results of forward speed of six cockroach robots changing with voltage. (B) Response results of left turning angle of six cockroach robots changing with voltage. (C) Response results of right turning angle of six cockroach robots changing with voltage.

## DISCUSSION

This paper designs a system for recording and analyzing the trajectory of the cockroach robot. The system builds an experimental platform for the cockroach robot to crawl freely. While the cockroach robot is remotely stimulated with appropriate electrical signal parameters, its movement trajectory can be recorded and stored in real-time, and post-processed by the trajectory data analysis module, which completes the quantitative analysis between stimulation parameters and movement behavior changes of the cockroach robot. The system improves the accuracy of behavioral studies of cockroach robots and simplifies the experimental process.

Tethering an insect to a trackball can affect the animal’s locomotor behavior ^[22,30]^; the insect is not able to move freely and its stimulus-response may be reduced. Instead, this system builds an infrared touchpad to provide an experimental platform for the cockroach robot to crawl. The larger screen size of the infrared touchpad and the higher sensitivity of recognition allows the cockroach to move freely and restore the insect’s locomotor instincts to the maximum.

The system simplifies the process of experimentation by changing from having the cockroach move on a trackball to having the cockroach move on a touchpad. The trackball system was changed from the original direct contact with two optical mice to record the insect’s trajectory ^[11,12]^, which had a mechanical resistance and could be inaccurate in recording the cockroach’s trajectory. On this basis, scientists proposed air-cushion trackballs to record insect trajectories ^[18]^, which effectively avoided the problem of mechanical resistance. Later, a virtual reality trackball system was proposed ^[29,30]^, which virtualized a natural environment to enable insects to go about their movements in a familiar environment and record their trajectories through trackballs. While recording insect trajectories with trackballs may have problems such as inertia, this paper constructs a system software module that can more intuitively see the trajectories of cockroaches’ movements through the projector’s interface projection on the infrared touchpad, and record and store the trajectories of cockroaches in real-time, which simplifies the recording method of trackballs, so there are no problems such as inertia and friction, and there is no need to tether the cockroaches, and the system is relatively accurate and can immediately display the cockroach’s movement trajectory on the infrared touchpad during its movement, which is the actual path the cockroach is walking on.

The system was designed with a data analysis module to store the movement trajectory of the cockroach, carry out analysis of the experimental data and observe the response under different stimulation parameters. Through the data analysis module, it was finally concluded that the forward speed, as well as the rotation angle of the cockroach robot, increased with the increase of voltage in the stimulation voltage range of 1-3 V, and the speed as well as the angle of the cockroach under the same stimulation voltage The findings include the changing pattern of the cockroach robot’s speed and angle of rotation in the same stimulation voltage.

The paper focuses on the overall design and experimental testing of the cockroach robot’s motion trajectory recording and analysis system. As the cockroach electrode implantation surgery cannot be precise and consistent every time, the experimental results of the system are prone to certain deviations. The size of the infrared touchpad is fixed and the free movement of the cockroach robot is irregular, so there is a problem of cockroaches crawling out of the experimental area during the experiment. Further research is still needed in the direction of recording and analyzing the movement trajectory of the cockroach robot, and the next steps need to be continued.

Increase the types of stimulus waveforms for the cockroach robot and explore the effects of different waveforms on the movement of the cockroach robot. Improve the algorithm for calculating the coordinate positioning and the forward and left-right steering of the cockroach robot, and optimize the algorithm for analyzing the trajectory data of the cockroach robot, so as to broaden the way of verifying the changes in the movement of the cockroach robot after stimulation. Optimize the functions and structure of the whole system to make the interface of the experimental operation form of each functional module concise and clear, and to maintain the sustainability and coherence of the operation of each functional module as far as possible.

## Acknowledgement

We are grateful to the participants of this study.

## Author contributions

Conceptualization, R.H. and H.Z.; methodology, R.H. and S.Y.; software, H.Z, R.Y., Z.Y.,Y.S and S.Y; hardware, H.Z., S.Y. and P.Z.; experimental test, R.H., S.Y and H.Z; writing—original draft preparation, H.Z.; writing—review and editing H.Z. and H.W.; project administration, R.H.; funding acquisition, J.Y. and H.W. All authors have read and agreed to the published version of the manuscript.

